# Dissecting the interactions of the ISG15-USP18-STAT2 inhibitory complex

**DOI:** 10.64898/2026.03.26.714284

**Authors:** Jessica C. Rowe, Yi Min Ng, Hannah Kiefer, Matthew Simmons, Marilyn Paul, Ramasubramanian Sundaramoorthy, David J. Hughes, Kirby N. Swatek

**Affiliations:** Medical Research Council Protein Phosphorylation and Ubiquitylation Unit, Faculty of Life Sciences, University of Dundee, Dundee, United Kingdom; Drug Discovery Unit, Faculty of Life Sciences, University of Dundee, Dundee, United Kingdom; Molecular Cell and Developmental Biology, Faculty of Life Sciences, University of Dundee, Dundee, United Kingdom; Biomedical Sciences Research Complex, School of Biology, University of St Andrews, St Andrews, United Kingdom

## Abstract

The suppression of type I interferon (IFN) signalling by the ISG15-USP18-STAT2 inhibitory complex (ISG15 IC) is an established regulatory mechanism of the antiviral response. However, a molecular understanding of how the ISG15 IC forms to suppress IFN signalling is still emerging. Here, we use AlphaFold modelling in conjunction with biochemical and biophysical approaches to elucidate the interactions of this multiprotein assembly. Our analysis identified a unique STAT2 binding loop (SBL) in USP18, which is critical for the USP18-STAT2 association. Further biochemical characterisation through site-directed mutagenesis confirmed the importance of residues within and surrounding the SBL, enabling the design of mutants with both increased and decreased binding affinities. Moreover, several USP18 and STAT2 patient mutations severely disrupted this interaction. Lastly, using influenza B virus (IBV) and Zika virus (ZIKV) proteins, we investigated the influence of these viral effector proteins on these interactions. Taken together, these results provide much-needed insights into a key aspect of IFN signalling control.

## Introduction

The detection of pathogens by the host elicits potent antiviral and inflammatory signalling responses to protect against the infection^1^. Antiviral cytokines called type I interferons (IFNs) are transcriptionally upregulated and secreted from virally infected cells^2^. Once secreted, IFNs act on neighbouring cells by binding to the heterodimeric IFN receptor (IFNAR1/IFNAR2), leading to the activation of receptor-associated tyrosine kinases JAK1 and TYK2 and subsequent phosphorylation and transcriptional activation of signal transducer and activator of transcription (STAT)1 and STAT2^3^. The outcome of these signalling events is the upregulation of hundreds of interferon-stimulated genes (ISGs).

The mechanisms of ISG-mediated antiviral immunity are diverse and dynamically regulated. While several ISGs are positive regulators of the antiviral response, including pattern recognition receptors (PRRs) which detect pathogens, and transcription factors that induce ISG expression (e.g. STAT2), negative regulators limit the degree and duration of the IFN response, often by directly antagonising signalling at the IFN receptor^2,4^. The suppressor of cytokine signalling (SOCS) protein family members, for example, inhibit JAK kinase activity, while protein tyrosine phosphatase 1B (PTP1B) and SRC homology 2 domain-containing protein tyrosine phosphatases (SHP-1 and SHP-2) target the IFN receptor-associated kinases and downstream STAT1 and STAT2 transcription factors^5^.

Another important mechanism of suppressing IFN signalling involves the ubiquitin-like protein interferon-stimulated gene 15 (ISG15). During the IFN response, ISG15 is one of the most highly upregulated ISGs^6,7^, and through its E1-E2-E3 enzymes becomes co-translationally attached to thousands of substrates^8–13^. As with other ubiquitin-like modifiers, ISGylation is a reversible modification. The deISGylase ubiquitin-specific protease 18 (USP18) specifically removes ISG15 from substrates ^14,15^. Intriguingly, independent of its catalytic activity, USP18 is a central negative regulator of type I IFN signalling^16^. Moreover, and in addition to its role as a modifier, unconjugated ISG15 (at least in human cells) binds to and stabilises USP18 and facilitates its negative regulatory activity^17^. It was recently shown that while a low affinity ISG15 mutant protected USP18 from degradation, wild-type ISG15 was required to drive negative regulation, further highlighting the importance of ISG15 in this pathway^18^. The mechanism of USP18-mediated IFN suppression is reported to occur through an association with the coiled-coil and DNA-binding domains of STAT2, which transport USP18 to the IFN receptor^16,19^. It has been suggested that the ISG15 IC inhibits IFN signalling by preventing IFNAR dimerization, possibly by displacing JAK1 from the IFN receptor, leading to a reduction in STAT1 and STAT2 phosphorylation^20,21^.

These negative regulators are crucial for maintaining a balanced immune response, and their dysregulation is associated with IFN-induced autoinflammation or type I interferonopathies. This association has been firmly established with the discovery of patients with ISG15 IC mutations. For example, loss of ISG15 in humans frequently results in symptoms consistent with mild type I interferonopathy, including intracranial calcification, seizures, skin lesions, and lung disease^17,22–27^. Remarkably, patients with ISG15 deficiency often reach adulthood and symptoms can be intermittent and resolve spontaneously^17,22,23^. Furthermore, these patients are not overtly prone to viral disease due to elevated IFN signalling^17,28,29^. In contrast, USP18 mutations are far more severe, with symptoms first appearing in the neonatal period and often proving fatal without therapeutic intervention^30–33^. Because STAT2 functions as both a positive and negative regulator of IFN signalling, diseases caused by monogenic STAT2 mutations span a broad clinical spectrum. Loss-of-function autosomal recessive mutations increase viral immunodeficiency^34^, whereas gain-of-function mutations that interfere with ISG15 IC formation phenocopy symptoms of USP18-deficient patients^35,36^. These clinical associations underscore the importance of deciphering the mechanisms and regulation of the ISG15 IC.

Therefore, we reconstituted the ISG15-USP18-STAT2 ternary complex and defined the interactions that underlie its assembly. Using a high-confidence AlphaFold model, we pinpoint the structural elements and amino acids required for USP18-STAT2 recruitment. Biochemical assays confirmed the molecular basis of these interactions, defining a unique loop within the palm domain of USP18 as a critical feature for binding the coiled-coil domain of STAT2. We further confirm that USP18 and STAT2 patient mutations disrupt these associations in quantitative binding assays. Our findings uncover a surprisingly intricate but delicate network of interactions and thereby provide new insights into how the ISG15 IC assembles to regulate IFN signalling.

## Results

### Assembly and characterisation of the ISG15 inhibitory complex

To characterise the ISG15-USP18-STAT2 complex, we purified each of these components individually or in complex with one another (**Fig. 1a, Supplementary Fig. 1**). Briefly, we co-expressed ISG15, Twin-Strep-tagged STAT2, and a His-tagged USP18 isoform (36-372 aa) using the biGBac baculovirus expression system^37,38^, and performed a tandem affinity purification (TAP) to isolate the intact complex. Based on Coomassie-stained SDS-PAGE gels the TAP-purified ISG15 IC subunits appear to be present at roughly similar levels with band intensity decreasing from high to low molecular weight proteins (**Fig. 1a**). However, upon analytical sizing the ternary complex dissociated with clear separation of STAT2 from the ISG15-USP18 association (**Supplementary Fig. 1b**). Based on this result, we then expressed and purified full-length STAT2 alone, as well as the human USP18-ISG15 complex.

**Figure 1.**
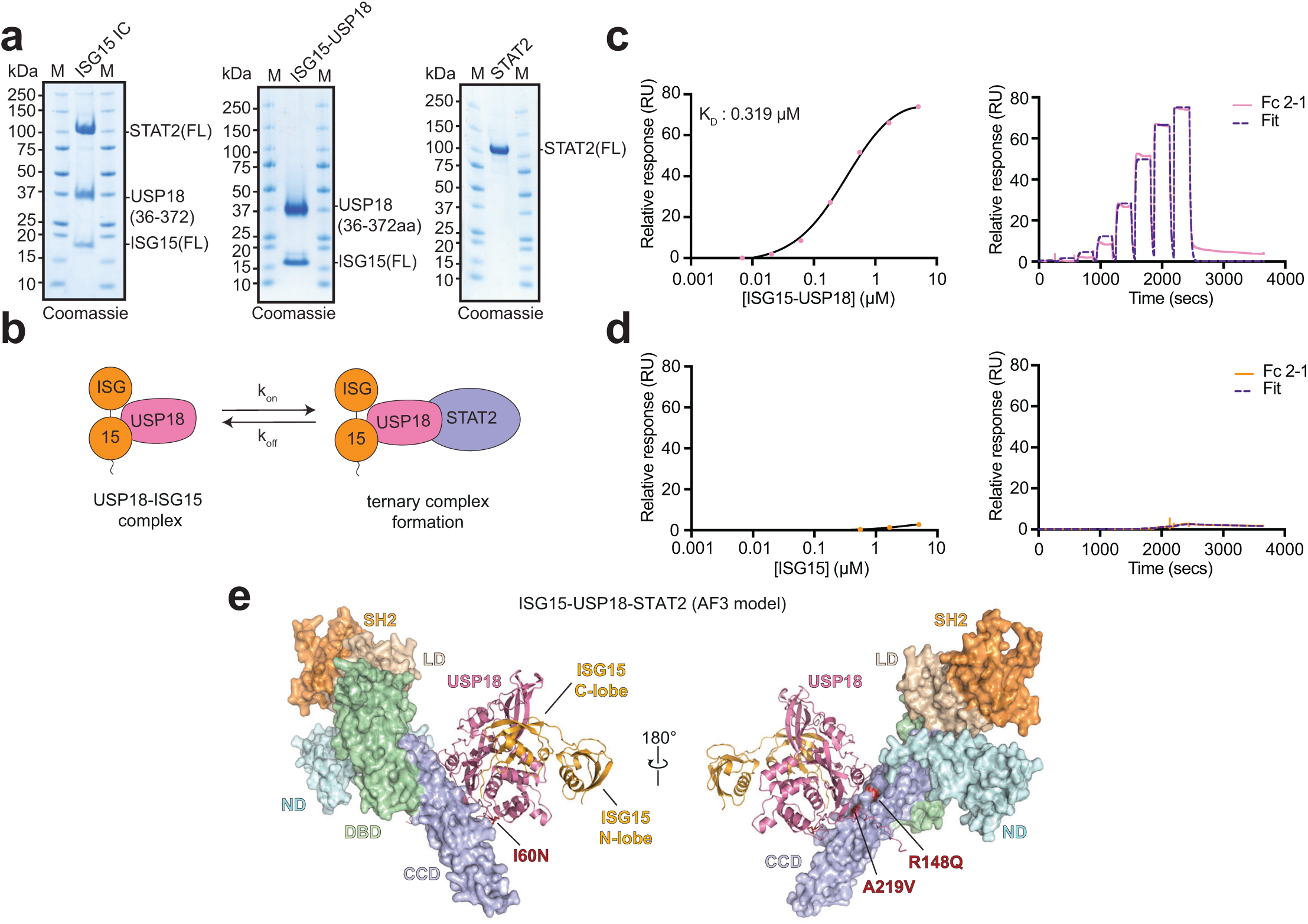
Assembly, binding, and structural modelling of the ISG15 inhibitory complex. **a**, Purification of the ISG15 inhibitory complex (IC) and its components. Purified proteins were visualised by Coomassie-stained SDS-PAGE gels (3 µg of each protein was loaded on the gel). **b,** Schematic depicting the SPR workflow. Co-expressed and purified ISG15-USP18 (analyte) was added to STAT2 (immobilised ligand) to determine the on and off rates of the interaction. **c, d,** Single-cycle SPR analysis of the interaction between STAT2 and ISG15-USP18 (**c**) or ISG15 only (**d**). A representative example of experiments performed independently in triplicate is shown (n=3). **e,** AlphaFold3 (AF) model of the ISG15 IC comprised of ISG15 (1-157 aa), USP18 (36-372 aa) and STAT2 (1-681 aa). STAT2 has been coloured according to its domains. ND: N-terminal domain; CCD: coiled-coil domain; DBD: DNA-binding domain; LD: Linker domain; SH2: SRC homology 2 domain.

With all components in hand, we next wanted to measure the binding affinity between STAT2 and ISG15-USP18. Using surface plasmon resonance (SPR), increasing concentrations of ISG15-USP18 were passed over the immobilised STAT2 ligand, allowing for determination of the steady-state binding affinity (**Fig. 1b, c**). This analysis revealed that human ISG15-USP18 bound to STAT2 with a binding constant (K_D_) of 0.350 (±0.048) µM (**Fig. 1c, Supplementary Fig. 2a-c**). Interestingly, this affinity is similar to that recently reported for the human ISG15-USP18 interaction^18^, suggesting equivalent binding strengths between each subunit. To investigate whether the USP18-STAT2 interaction is direct and not a result of ISG15 binding, we assessed the interaction between STAT2 and ISG15 in biochemical assays. In these assays no binding was observed indicating that USP18 or the ISG15-USP18 complex is responsible for the STAT2 interaction (**Fig. 1d, Supplementary Fig. 2d-f**). This result is in agreement with a previous yeast-two-hybrid study between human USP18 and STAT2^19^.

To visualise the ISG15 IC we next performed cryo-EM analysis of the TAP-purified complex. Cryo-EM revealed clear two-dimensional (2D) and three-dimensional (3D) classes and produced a low-resolution map which enabled the placement of each subunit within the cryo-EM density, providing evidence the complex exists as a heterotrimer (**Supplementary Fig. 2 g-i, Fig. 1a**). However, due to particle orientation bias it was not possible to generate a high-resolution 3D reconstruction of the complex. Therefore, we turned to AlphaFold3 (hereafter referred to as AlphaFold) to model the complex. Rewardingly, AlphaFold produced a high confidence model (pTM:0.7, iPTM:0.69) that resembled our low-resolution cryo-EM data (**Fig. 1e**)^39^. Moreover, the AlphaFold model supported our binding data that USP18 is the STAT2 contact site (**Fig. 1c, d**). Together, the model revealed several important features of the ISG15 IC, including that the USP18-STAT2 interaction occurs through the USP18 palm domain and STAT2 coiled-coil domain, ISG15 only interacts with USP18, and sites of known patient mutations are located within the USP18-STAT2 interface (**Fig.1e**).

### Identification of a unique STAT2 binding loop in USP18

To further understand the molecular basis of the USP18-STAT2 interaction we combined phylogenetic, structural, and biochemical analyses to define the specificity determinants of this association. Phylogenetic comparison revealed USP30 to be the most closely related USP family member to USP18 (**Fig. 2a**). Sequence alignment and conservation analysis of USP18 revealed a unique 10 amino acid insertion, which we refer to as the STAT2 binding loop (SBL) (**Fig. 2b, Supplementary Fig. 3**). The location of the SBL is on the back of the palm domain, while ISG15 is grasped between the thumb, fingers, and front of the palm domain (**Fig. 2c**). Overlaying a USP30 crystal structure onto the ISG15-USP18 AlphaFold model revealed several surface-exposed hydrophobic and negatively charged residues, which were predicted to be important for STAT2 binding (**Fig. 2d**)^40^. Indeed, as predicted, USP30 was void of STAT2-binding activity in fluorescent polarisation (FP) binding assays despite increasing protein concentrations, in stark contrast to ISG15-USP18 (**Fig. 2e**). This analysis suggests the SBL could be a key determinant of STAT2 recognition by USP18.

**Figure 2.**
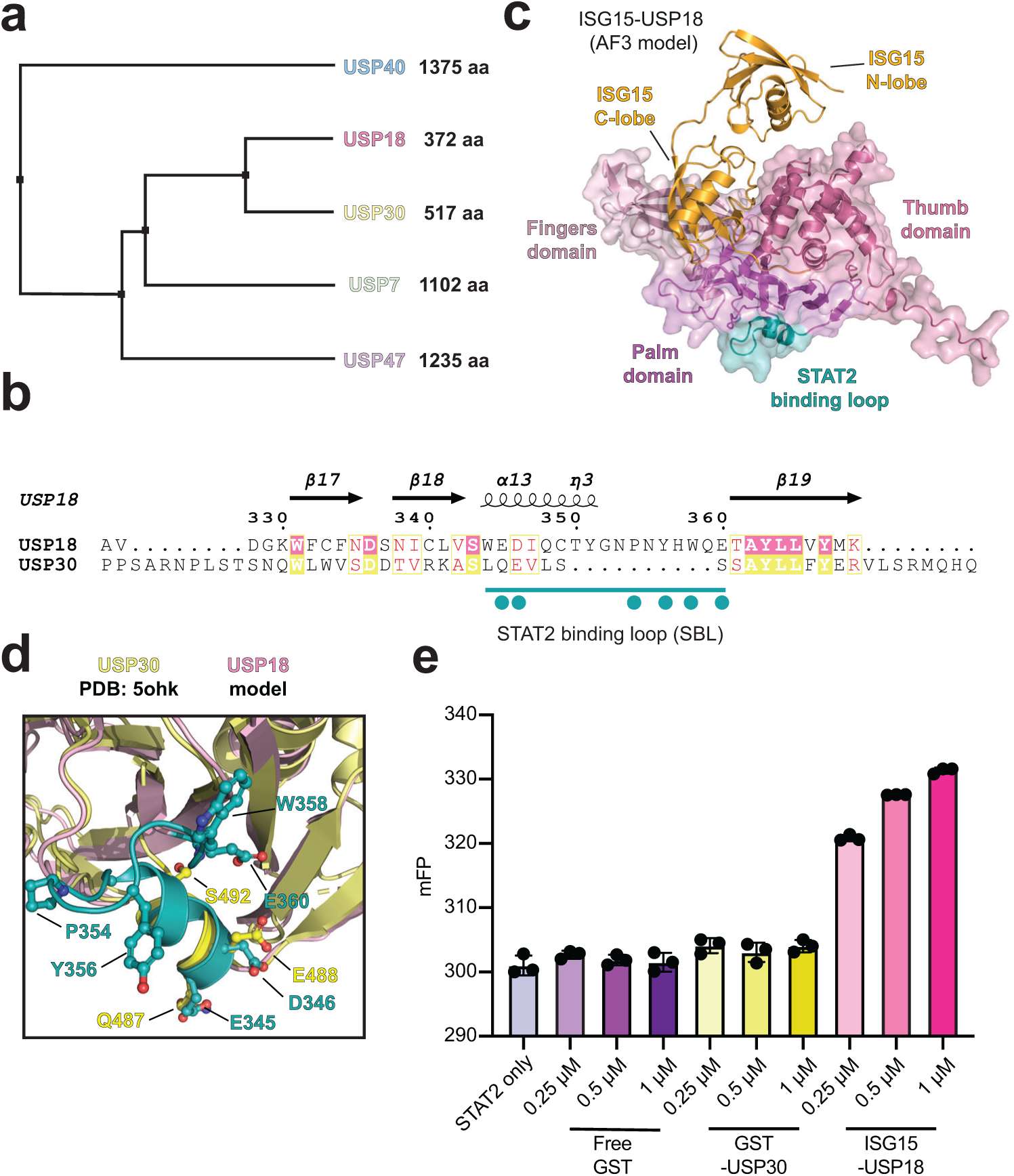
USP18 contains a distinct STAT2 binding loop. **a,** Phylogenetic tree of the USP clade containing USP18. The phylogenetic tree was generated using sequence alignment in Jalview and is drawn to scale, with branch lengths reflecting relative evolutionary distances^56^. **b,** Multisequence alignment of USP18 and USP30. T-Coffee and ESPript were used to perform the sequence alignments^57,58^. The teal line indicates the SBL and the teal circles denote surface exposed charged and hydrophobic residues. **c,** AlphaFold model of human USP18 bound to ISG15 with the USP domains and SBL highlighted. Thumb domain: 1-168 aa, Finger domain: 169-191 aa and 210-247 aa, Palm domain: 192-209 aa and 248-372 aa, SBL: 343-360 aa. **d,** Structural overlay of the USP18 AlphaFold model with the USP30 crystal structure (PDB:5OHK) highlighting the absence of the SBL in USP30^40^. **e,** Fluorescence polarisation (FP) binding assay comparing ISG15-USP18 versus USP30 STAT2 binding. Free-GST was used as a negative control. The bar graph is a representative example of three independent experiments performed in technical triplicate (n =3).

### Analysis of USP18 mutants on the STAT2 interaction

With binding assays to study the ISG15-USP18 and STAT2 interaction established, we next tested a comprehensive panel of USP18 mutants for defects in STAT2 binding. Using the previously mentioned high confidence AlphaFold model as reference, we were able to predict and design several site-specific mutants likely to disrupt ternary complex formation. These residues were primarily located in the SBL of the palm domain, with additional sites in the thumb domain (**Fig. 2c**, **Fig. 3a**). Hydrophobic residues were mutated to alanine (V311A, Y356A, W358A and Y356A/W358A), while residues involved in ionic interactions were exchanged for amino acids of the opposite charge (R119D, E345K, D346K, E360K, and D346K/E360K). Importantly, E345 and D346 were predicted to interact with R148 of STAT2, the site of a gain-of-function mutation found in patients with interferonopathies^36^. The catalytically inactive C64A and patient-derived USP18 I60N mutants were also included. In total, eleven different ISG15-USP18 mutants were purified (**Fig. 3b**).

**Figure 3.**
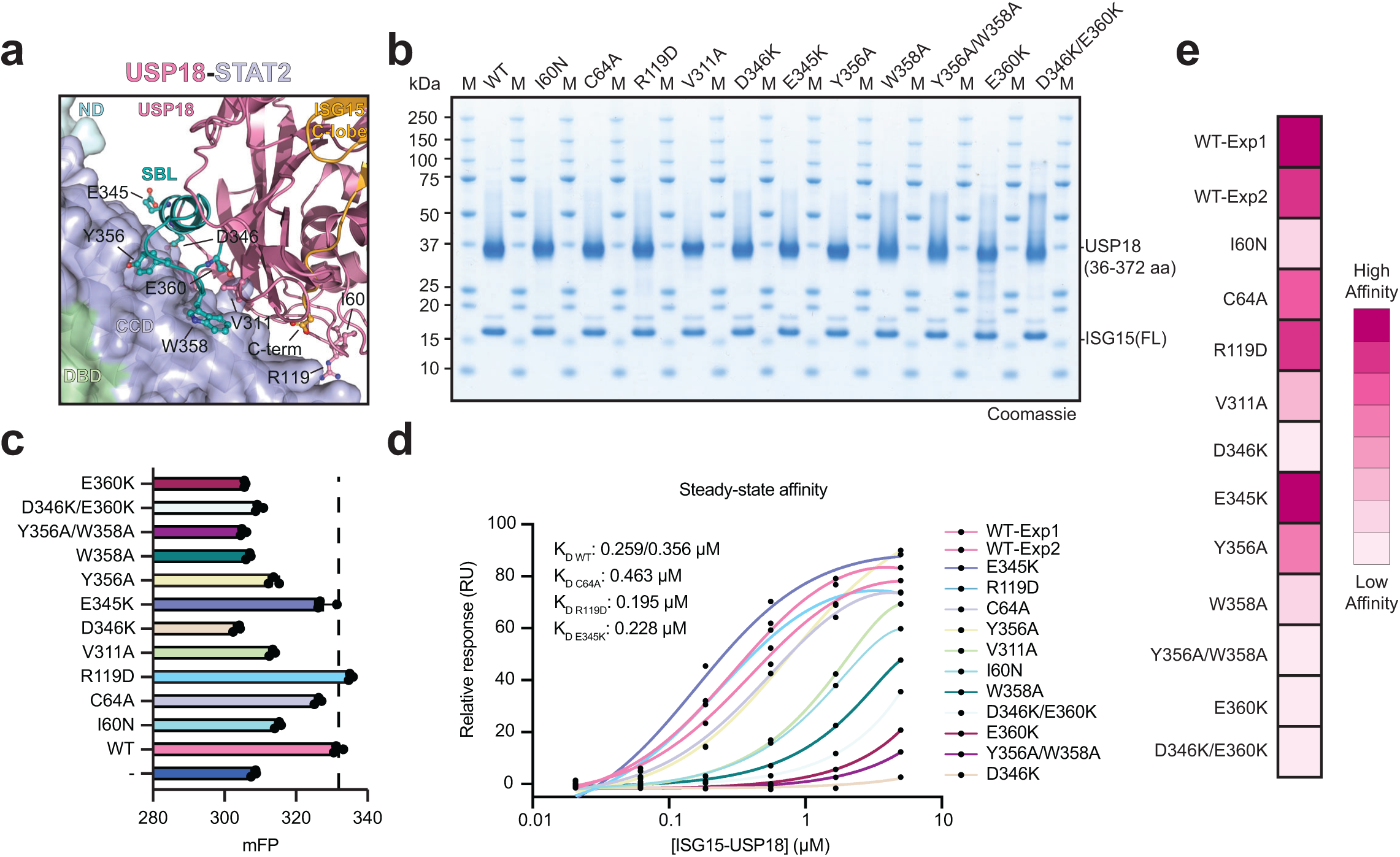
Binding of ISG15-USP18 mutants to STAT2. **a**, Close-up view of the USP18-STAT2 interaction site based on the AlphaFold model (also see Fig. 1e). Highlighted USP18 residues are predicted to interact with the coiled-coil domain (CCD) of STAT2 and were selected for further characterisation by site-directed mutagenesis binding studies. **b,** Purification of the ISG15-USP18 mutants. Approximately 3 µg of each ISG15-USP18 binary complex was separated by SDS-PAGE and visualised with Coomassie stain. **c,** Fluorescence polarisation (FP) binding assays between ISG15-USP18 mutants and FlAsH-tagged STAT2. The bar graph is a representative example of three independent experiments performed in technical triplicate (n=3). **d,** Surface plasmon resonance binding assays between the ISG15-USP18 mutants and STAT2. The binding curve plots are a representative example of independent experiments performed in triplicate (n=3) **e,** Binding affinity heatmap of the ISG15-USP18 mutants. Boxes are coloured according to normalised median RU values (i.e., the RU value closest to wild-type K_D_). Colour intensity reflects binding from low affinity (light pink) to high affinity (dark pink).

Initial screening of our panel of USP18 mutants in qualitative FP binding assays against FlAsH-tagged STAT2 revealed that seven out of the nine USP18 structural mutants exhibited reduced STAT2 binding in comparison to controls (**Fig. 3c**, **Supplementary Fig. 4n**). The only two exceptions were the R119D and E345K mutations, which displayed binding levels comparable to wild-type USP18. Additionally, and in agreement with previous reports^18,19,41^, the catalytically inactive USP18 bound STAT2 at comparable levels to wild-type (**Fig. 3c**). Thus, our biophysical assays confirmed several structural determinants of the USP18-STAT2 interaction.

To quantitatively assess the impact of these USP18 mutants on ISG15 IC formation in a concentration-dependent manner, we next turned to SPR binding assays. Consistent with the results of the FP assays, the same mutants also showed reduced binding in these experiments (**Fig. 3d, e, Supplementary Fig. 4a-m, Supplementary Table 1**). Notably, SPR allowed us to confidently rank the magnitude of these defects. Five mutations scattered throughout the USP18 SBL domain (D346K, D346K/E360K, Y356A/W358A, W358A, E360K) displayed significantly weaker SPR binding signals compared to the clinically relevant I60N mutation (**Fig. 3a, d, e**), which is well-known to disrupt STAT2 binding and IFN regulation^31,42^. The most defective mutant was D346K, which is highly conserved across species and structurally positioned between two positively charged STAT2 residues (R148, R223) (**Supplementary Fig. 3, Fig. 3a**). Interestingly, the hydrophobic mutants also proved essential; while W358A significantly impaired binding, the Y356A/W358A double mutation nearly abolished the interaction. This additive effect indicates that these residues form a coordinated hydrophobic interface which is required to stabilise the complex. Conversely, the E345K mutation did not interfere with STAT2 binding, suggesting this position is less critical for the interaction.

### The impact of STAT2 mutations on ISG15 inhibitory complex formation

Since patient-derived STAT2 mutations at positions (A219, R148) are well known to interfere with the regulation of the ISG15 IC^35,36,43^, we sought to assess the extent to which these clinically relevant mutants disrupt complex formation. Concurrently, we investigated whether the USP18-STAT2 interface could be rationally engineered to enhance binding. Examination of the AlphaFold USP18-STAT2 interface revealed a hydrophobic contact site between USP18 W358 and two STAT2 residues within the coil-coiled domain (L299, A302) (**Fig. 3a**). We hypothesised that swapping these residues with a more hydrophobic amino acid might promote the USP18-STAT2 interaction (L299W or A302W) (**Fig. 4a, left**). Along with the hydrophobic mutants, we also expressed and purified the R148Q and A219V STAT2 patient mutations and tested their ability to bind ISG15-USP18 by quantitative SPR (**Fig. 4a, right, b, Supplementary Fig. 5, Supplementary Table 2**).

**Figure 4.**
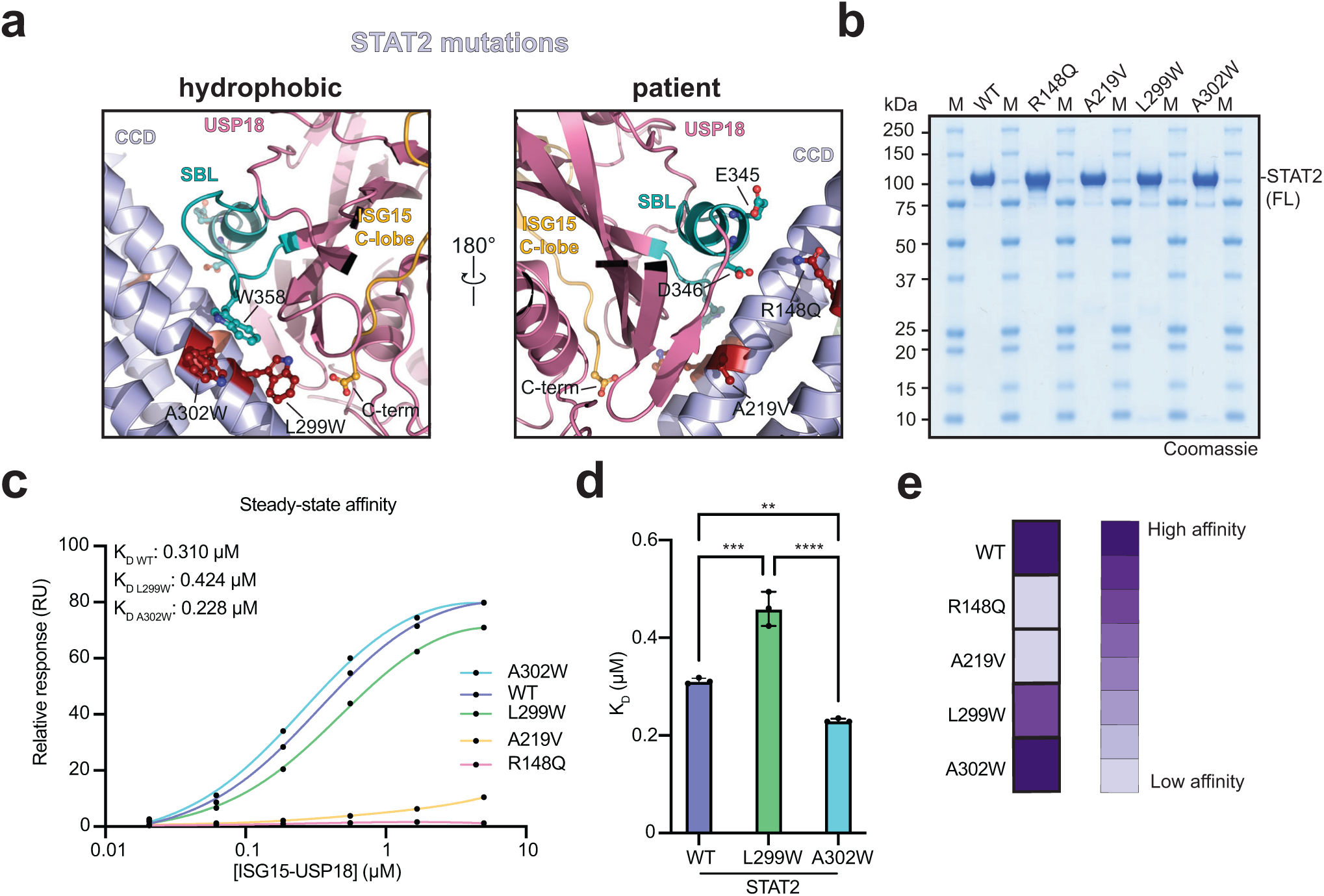
Binding of STAT2 mutants to the ISG15-USP18 complex. **a**, Close-up view of the ISG15 IC AlphaFold= model highlighting the interaction sites between the coiled-coil domain (CCD) of STAT2 and USP18. STAT2 residues highlighted in the left panel were selected for mutagenesis as shown. STAT2 residues highlighted in the right panel are identified patient mutations. **b,** Coomassie-stained SDS-PAGE gel of the purified STAT2 mutants. Approximately 3 µg of each STAT2 was loaded on the gel. **c,** Surface plasmon resonance binding studies between STAT2 mutants and the ISG15-USP18 complex. The binding curve plots are a representative example of independent experiments performed in triplicate (n=3). **d,** Binding affinities of STAT2 mutants. The mean K_D_ values for STAT2 and ISG15-USP18 binding are shown with error bars representing the standard deviation from independent experiments performed in triplicate (n=3). Statistical significance was assessed using one-way ANOVA and Tukey multiple comparisons test, **p<0.002, ***p<0.0002, ****p<0.0001. **e,** Binding affinity heatmap of STAT2 mutants. Boxes are coloured according to normalised median RU values (i.e., RUs closest to wild-type K_D_). Colour intensity reflects binding from low affinity (light purple) to high affinity (dark purple).

In agreement with previous co-immunoprecipitation (Co-IP) studies^35^, the A219V patient mutation was defective in USP18 binding (**Fig. 4c**). However, R148Q, which was not previously shown to inhibit USP18 binding in co-IP assays^36^, did not bind USP18 (**Fig. 4c**). The complete loss of binding is consistent with our earlier observation that mutating its interacting partner on USP18 (D346K) severely impairs complex formation, further confirming the importance of the R148-D346 electrostatic bridge. This discrepancy in literature likely reflects different experimental approaches; however, it is worth noting that another STAT2 R148 patient mutation (R148W) had reduced USP18 binding in co-IP assays^43^.

Beyond the clinical variants, the hydrophobic STAT2 mutations both positively and negatively impacted USP18 binding. As shown in the model, STAT2 A302W is directly below USP18 W358 and well positioned to form a hydrophobic interface (**Fig. 4a, left**). Indeed, the binding affinity of the STAT2 A302W mutant was slightly but significantly improved compared to wild-type STAT2 (K_D_ STAT2_WT_- 0.311 (±0.061) µM vs K_D_ STAT2_A302W_ – 0.230 (±0.004) µM) (**Fig. 4c, d, Supplementary Table 2**). The STAT2 L299W mutation, however, is nearly a full α-helical turn away from W358 and did not improve USP18-STAT2 binding but instead weakened the interaction (K_D_ STAT2_WT_- 0.311 (±0.061) µM vs K_D_ STAT2_L299W_ – 0.459 (±0.035) µM) (**Fig. 4**). Together, these results highlight that STAT2 patient mutations display little to no USP18 binding, while targeted optimisation of the hydrophobic interface with a site-specific A302W STAT2 mutation enhanced ternary complex formation.

### The influence of STAT2 on USP18’s catalytic activity

Since USP18 has two primary functional roles as a deISGylase and a suppressor of IFN signalling, we wanted to test if STAT2 influenced USP18 catalytic activity. Using proISG15 as a substrate, which contains eight amino acids past the C-terminal GlyGly residues, time-dependent cleavage assays revealed that the addition of STAT2 resulted in no change in USP18 cleavage efficiency (**Fig. 5a**). To further validate this result, increasing concentrations of STAT2 were added to the USP18 cleavage assays. Consistently, we observed that STAT2 had a minimal impact on proISG15 cleavage levels even at the highest concentration tested (**Fig. 5b**). Moving forward, it will be interesting to test if STAT2-binding impacts USP18’s ability to cleave ISGylated substrates, which will further characterise its substrate preference and mechanism of forming the ISG15-USP18 complex.

**Figure 5.**
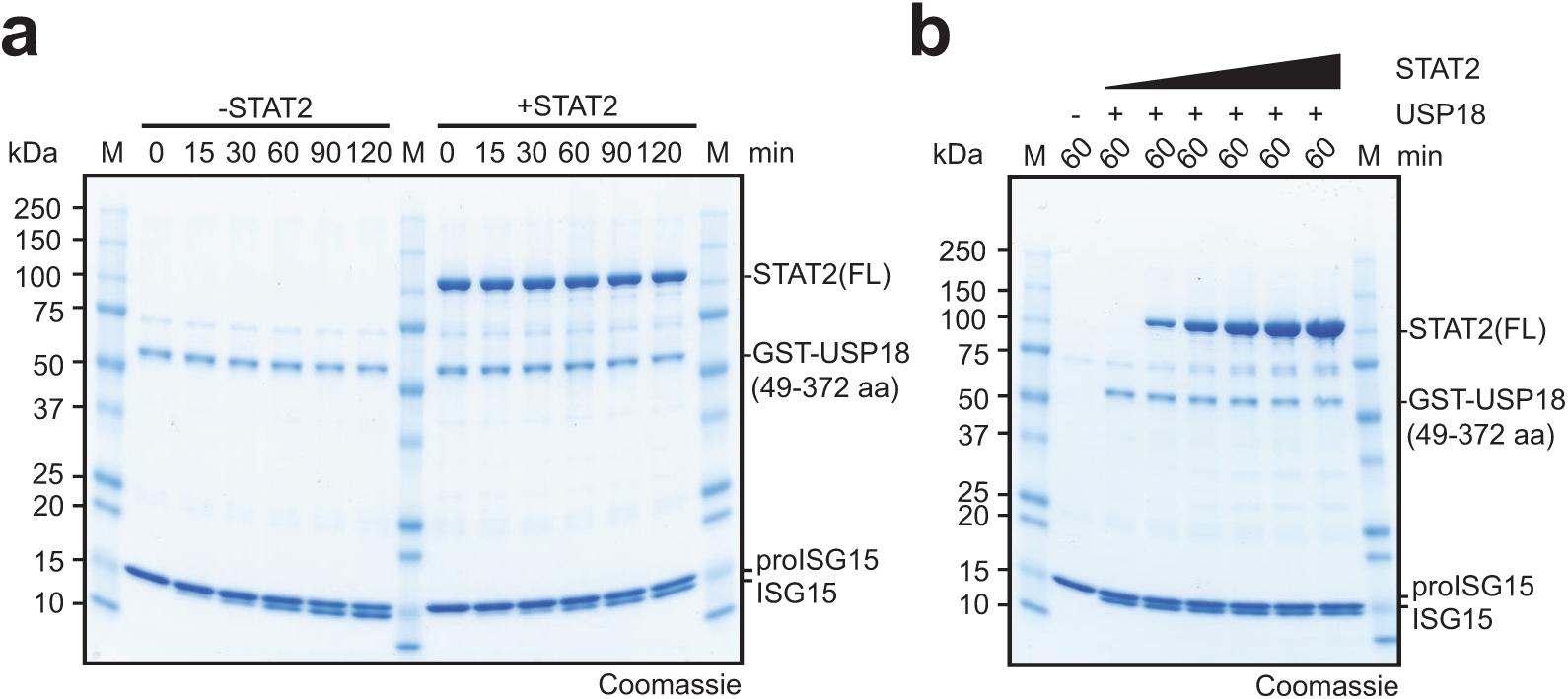
The impact of STAT2 on USP18 catalytic activity. **a,** Time-dependent USP18 cleavage assays with proISG15, minus and plus STAT2. The cleavage of proISG15 into mature ISG15, as indicated by the ISG15 mass shift, was visualised by Coomassie-stained SDS-PAGE gels. A representative example of experiments performed independently in triplicate is shown **b,** USP18 cleavage assays with proISG15 and increasing concentrations of STAT2. Reactions were visualised as in **a**. A representative example of experiments performed independently in triplicate is shown.

### The effects of viral proteins on ISG15 IC formation

Due to the antiviral activity of the IFN system, many viruses have evolved mechanisms to evade or suppress this immune response. For example, ISG15 is targeted by several viruses, including coronaviruses, nairoviruses, and influenza B virus^44,45^. Moreover, viral effector proteins from flaviviruses and paramyxoviruses can induce the proteasomal degradation of STAT2^46,47^. We previously used viral effector proteins to understand the mechanisms of ISG15 E1 activity^48^, and were curious if we could similarly use this strategy to further characterise the assembly of the ISG15 IC. Using fluorescently labelled STAT2, the non-structural protein 5 of Zika virus (NS5), and the non-structural protein 1 of influenza B virus (NS1B), we tested the impact of these effector proteins on ISG15 IC formation by real-time FP binding assays (**Fig. 6**)^49^. As expected, wild-type ISG15-USP18 bound STAT2, as indicated by a robust increase in FP signal, whereas a STAT2 binding defective USP18 mutant (Y356A/W358A) produced a modest increase (**Fig. 6b**). The addition of NS5 to both these reactions resulted in a further increase in FP signal, which was higher than STAT2-NS5 alone, suggesting formation of a higher order assembly (**Fig. 6b**). This result was somewhat unexpected considering a cryo-EM structure of NS5 in complex with STAT2 showed the methyltransferase domain of NS5 binds to the same location as USP18^50^. However, our data indicate that USP18 and NS5 can simultaneously bind STAT2. Like NS5, NS1B caused a further increase in FP signal when added to ISG15-USP18-STAT2, suggesting formation of a quaternary complex (**Fig. 6c, left**). Moreover, the ISG15-binding defective NS1B mutant (NS1B AA) did not bind (**Fig. 6c, right**). This was as expected since NS1B binds to the N-terminal ubiquitin-like domain and linker region of ISG15, which based on the AlphaFold model do not contribute to complex formation (**Fig. 1e**). These results indicate that both viral proteins can exist in complex with the ISG15 IC (**Fig 6d**), suggesting that studying how these viral effector proteins influence the regulation of this complex could be an interesting avenue for future research.

**Figure 6.**
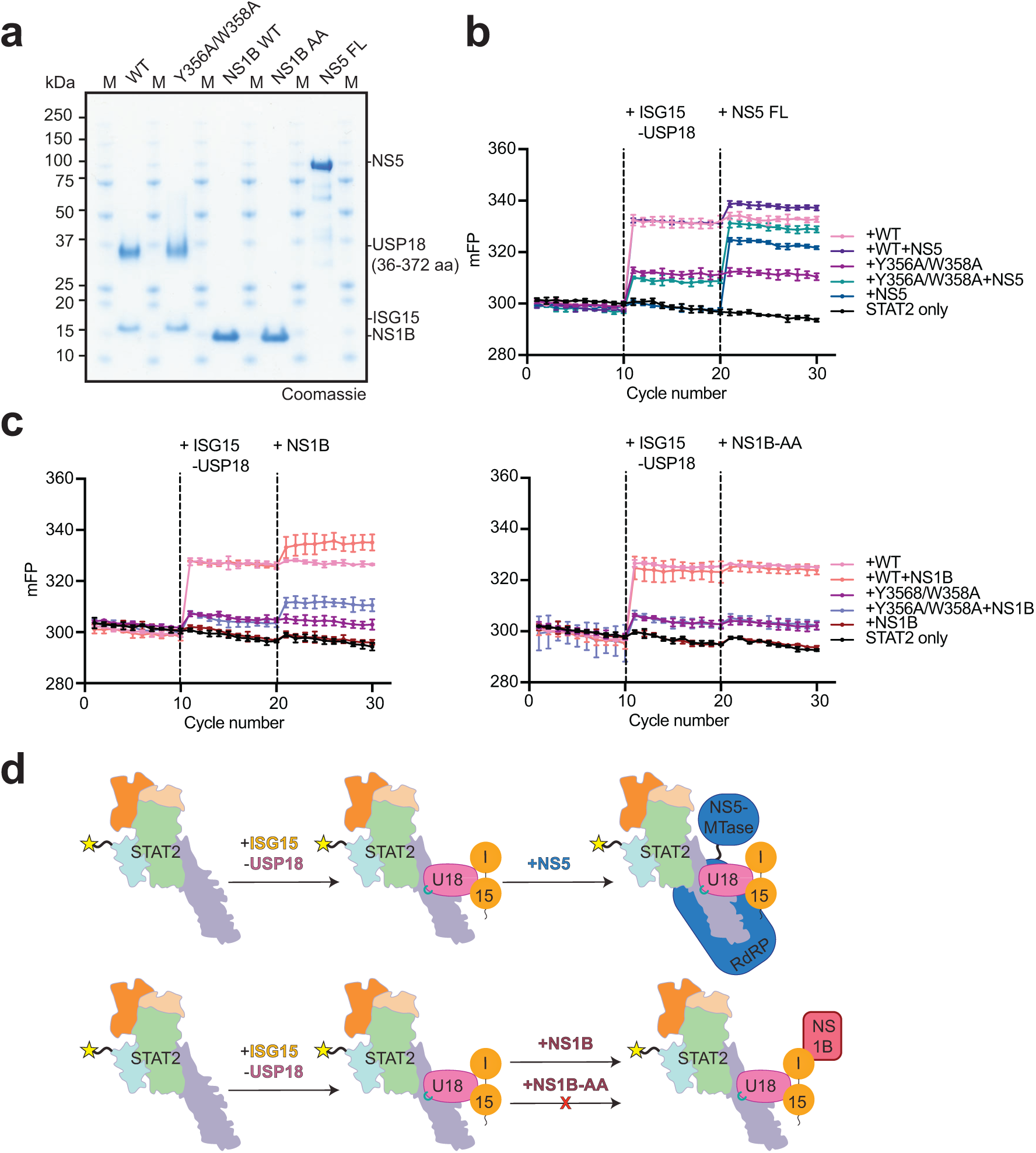
Effects of viral proteins on ISG15 inhibitory complex formation. **a,** Purified proteins used for binding studies were visualised by Coomassie-stained SDS-PAGE gels. Approximately, 3 micrograms of each protein was loaded on the gel. **b, c,** Fluorescence Polarisation (FP) binding assays of ISG15 IC formation in the presence of the viral proteins Zika virus NS5 (**b**) and influenza B virus wild-type NS1B or the ISG15 non-binding mutant, NS1B AA (**c**). Representative examples of experiments performed independently in triplicate is shown (n=3). The USP18 Y356A/W358A mutant was used as a control for STAT2 binding. **d,** Schematic model explaining the observed FP changes in **b** and **c**. Both NS5 and NS1B bind to the ISG15 IC, suggesting formation of a quaternary complex.

## Discussion

The mechanisms by which ISG15, USP18, and STAT2 assemble to suppress IFN signalling have remained unclear. Here, we explore this area by defining the surfaces that mediate the assembly of this complex using biochemical and biophysical approaches. Our comprehensive analysis identified several key residues within specific USP18 and STAT2 domains. When mutated, these residues disrupted ternary complex formation, revealing key determinants that facilitate assembly. The identification of the SBL is in agreement with a recent preprint reporting a cryo-EM structure of the ISG15 IC that was determined by trapping the complex with the SpyCatcher-SpyTag system^51^. On the other hand, their structure revealed a couple of additional contact sites, including the USP18 I60 binding site and the identification of a unique interaction within the USP18 thumb domain facilitated by a structural rearrangement of the STAT2 coil-coiled domain^51^. Another notable difference between our studies is that Huynh et al. detect a STAT2-dependent inhibition of USP18 activity which was not observed in our time- and concentration-dependent cleavage assays. Together, these reports address a fundamental question in the ISG15 field of how the ISG15 IC assembles at a molecular level, while also highlighting several important knowledge gaps in our current understanding of the downstream regulation of this pathway.

The assembly of the ISG15 IC is only the first step in its role as a suppressor of the IFN response. The signalling events that drive its localisation to the IFN receptor, and the molecular mechanisms by which it displaces JAK1 from IFNAR2, remain poorly understood. Furthermore, once bound to the receptor, how the ISG15 IC is removed or degraded to allow IFN signalling to resume has yet to be established. Building on our study and the recent preprint, it may soon be possible to visualise and biochemically characterise these higher-order assemblies. Since the ternary complex is inherently unstable, as shown by its dissociation in size-exclusion chromatography and the need to covalently trap the complex for high-resolution cryo-EM structural determination, it seems plausible that the reconstitution of these larger multiprotein complexes may facilitate the stabilisation of the ISG15 IC subcomplex.

The formation of viral effector-containing ISG15 IC assemblies may also regulate this complex during infection. Our FP binding assays clearly show that Zika virus NS5 and Influenza B virus NS1B interact with this complex (**Fig. 6**). In agreement with this finding, NS5 has recently been shown to bind STAT2 and USP18 in cells^41^. However, increasing concentrations of USP18 were reported to outcompete NS5 for STAT2 binding^41^. Since NS5 suppresses IFN signalling by promoting the proteasomal degradation of STAT2^52,53^, a mechanism was proposed in which ISG15-USP18 protect STAT2 from degradation^41^. This ISG15-USP18 protection mechanism may be short-lived due to increasing NS5 concentrations during viral infection but could still provide the host enough time to mount an effective immune response. Nevertheless, this further expands the regulatory roles of the ISG15 system, suggesting that the ISG15-USP18 complex can also potentiate the IFN response during Zika virus infection. Future studies are required to understand the biological implications of the NS1B interaction. However, our data suggest that this interaction, as well as other viral proteins that bind ISG15 IC subunits, warrant further investigation.

Due to its role as a key regulator of the IFN response, the ISG15 IC has emerged as a promising therapeutic target for diseases including cancer, immune disorders, and viral infections^54,55^. This study indicates that the ISG15-USP18-STAT2 interaction can be fine-tuned through site-specific mutations. Whether these differences in affinity lead to a proportional change in the magnitude of IFN signalling remains to be established. Encouragingly, many of the USP18 mutations designed here bind with lower affinity than the USP18 I60N patient variant, which led to elevated IFN signalling in cells and humans^18,31^. Moreover, the STAT2 A302W mutant demonstrates that this complex can be stabilised. Future studies should therefore define the degree and duration to which these mutants modulate IFN signalling in cells, which would serve as proof-of-principle that the mechanistic insights gained here can be leveraged to tune the IFN response. If successful, this could reveal discrete ‘hotspots’ that could be targeted by small molecules to selectively enhance or dampen IFN signalling. Together, the ability to reconstitute this pathway and quantitatively monitor these interactions, as reported here, provides a foundation for developing small molecules that target the ISG15 IC, thereby facilitating a deeper understanding of the therapeutic potential of modulating this key suppressor of innate immune signalling.

## Acknowledgments

We thank the members of the Swatek Lab and MRC PPU Reagents and Services for meaningful discussions and reagents. This study was supported by the Medical Research Council (MC_UU_00018/10; MC_UU_00038/8) and Royal Society (RGS_R2_222185) to K.N.S. K.N.S is a Lister Institute Prize Fellow.

## Author contribution

J.C.R. designed assays, performed experiments, interpreted results, and edited the manuscript. Y.M.N., H.K., and M.S. performed experiments and prepared reagents. M.P. assisted in designing and interpreting SPR assays. R.S. processed the cryo-EM data. D.J.H. assisted in the designing of experiments. K.N.S conceived and designed the study, performed experiments, interpreted results, wrote the manuscript and acquired funding.

## Conflict of Interest Statement

The authors declare no conflicting interest.

---

**Supplementary Figure 1.**
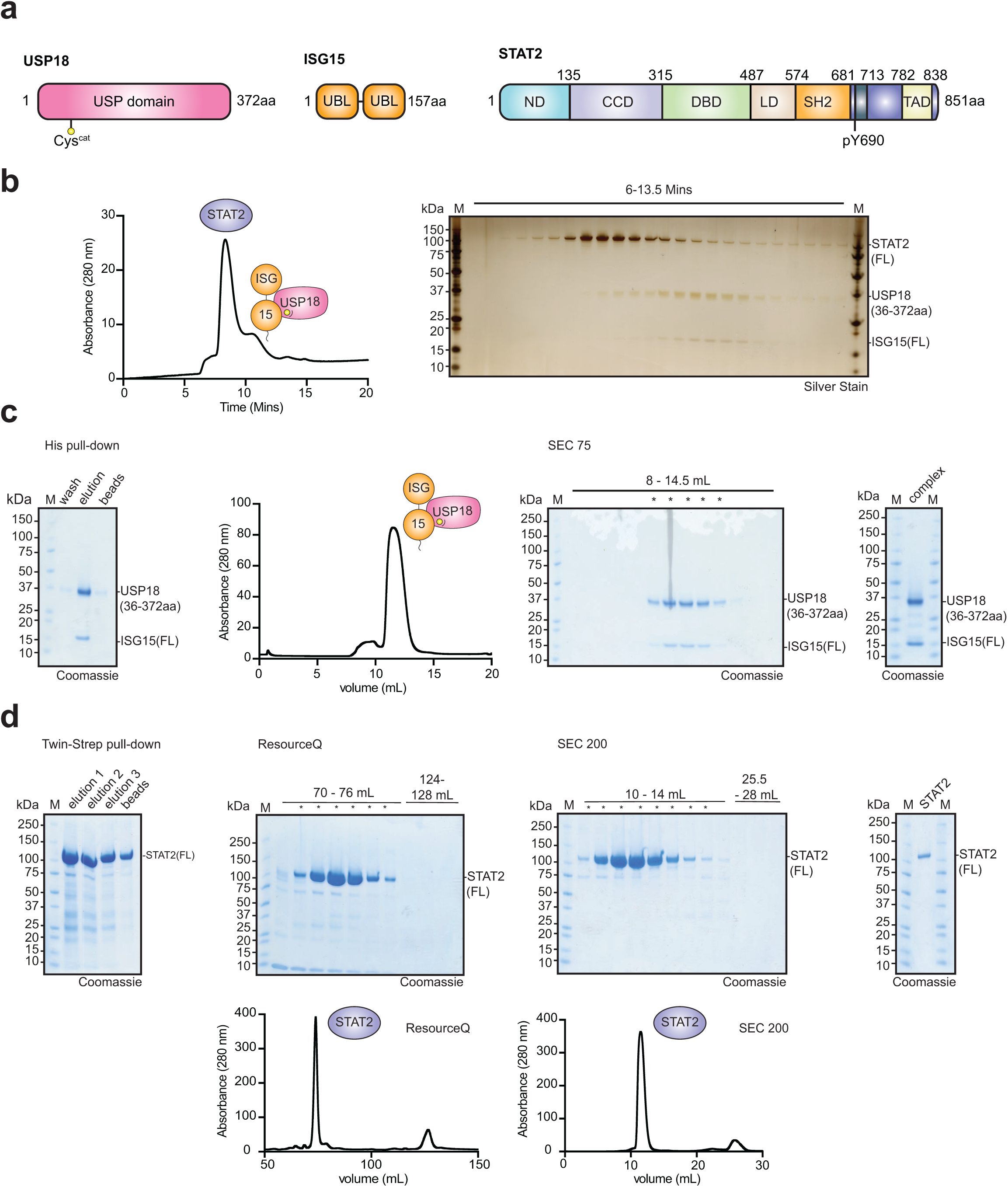
Purification of the ISG15 Inhibitory complex components. **a**, Domain schematic of the ISG15 inhibitory complex components STAT2, USP18 and ISG15. ND: N-terminal domain; CCD: Coiled-coil domain; DBD: DNA-binding domain; LD: Linker domain; SH2: SRC homology 2 domain; TAD: Transcriptional activation domain; USP: Ubiquitin-specific peptidase domain; UBL: Ubiquitin-like domain. **b,** High-performance liquid chromatography (HPLC) of the ISG15 inhibitory complex. Fractions were loaded onto an SDS-PAGE gel and visualised by silver stain. **c,** Purification of the wild-type ISG15-USP18 complex. Left to right: SDS-PAGE gel of the His-tag enrichment step; analytical size exclusion chromatography (SEC) of the ISG15-USP18 binary complex; fractions from the SEC run were visualised with Coomassie staining and those marked with an asterisk were combined and concentrated; Approximately 3 µg of the purified complex was loaded on the SDS-PAGE gel and visualised with Coomassie stain. **d,** Purification of wild-type STAT2. Left to right: SDS-PAGE of the Twin-Strep enrichment step; Ion exchange chromatography using a ResourceQ column – fractions marked with an asterisk in the Coomassie-stained SDS-PAGE gel were pooled and concentrated; Analytical SEC of STAT2 – fractions marked with an asterisk in the Coomassie-stained SDS-PAGE gel were pooled and concentrated; Approximately 2 µg of wild-type STAT2 was loaded on the SDS-PAGE gel and visualised with Coomassie stain.

**Supplementary Figure 2.**
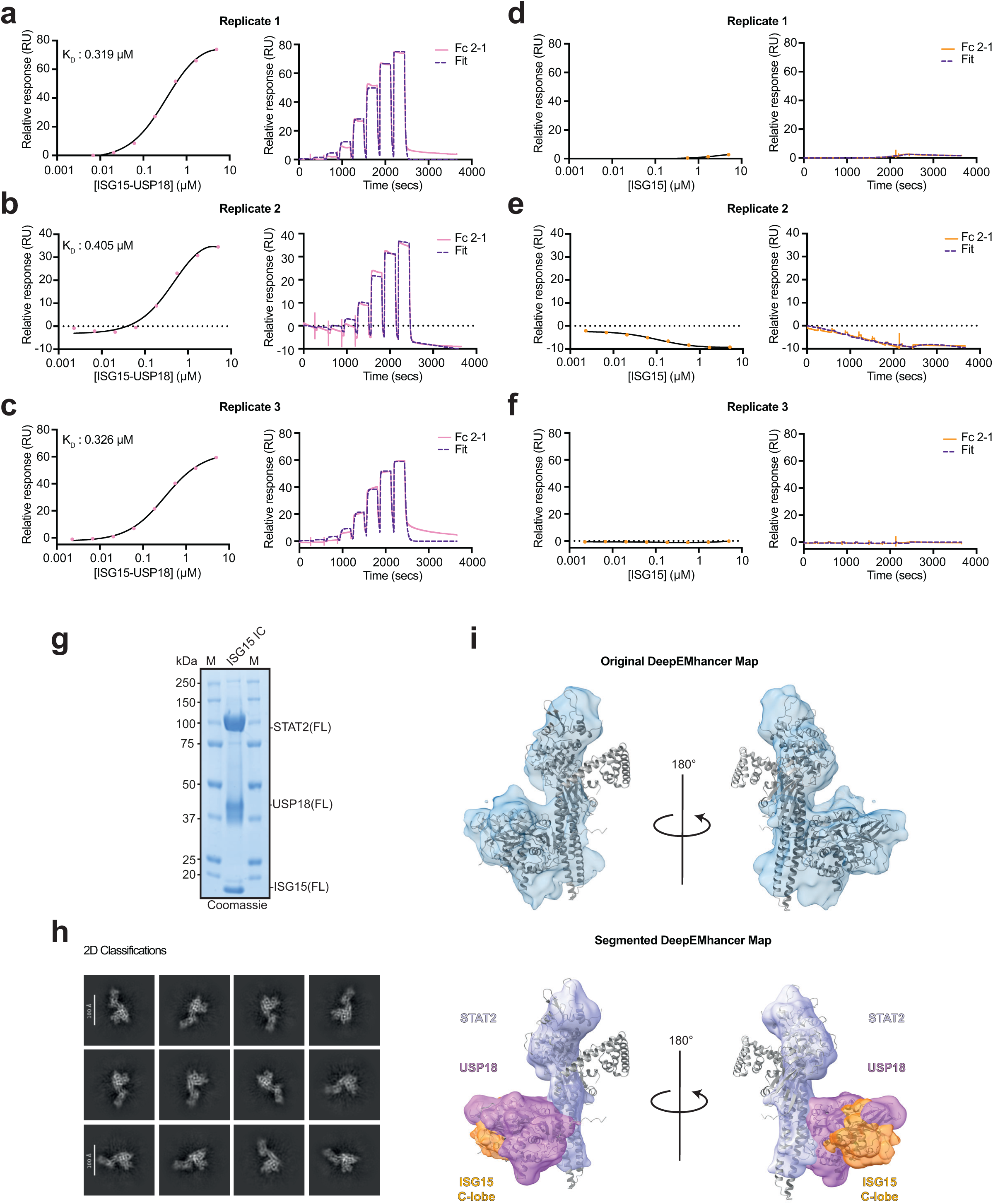
SPR binding assays and cryo-EM analysis of the ISG15 IC. **a-c**, Independent experiments performed in triplicate are shown for ISG15-USP18 binding against STAT2 (n=3). **d-f**, Independent experiments performed in triplicate are shown for ISG15 binding against STAT2 (n=3). For all experiments, wild-type STAT2 was immobilised onto a Strep-Tactin coupled sensor chip and association and dissociation rates of either ISG15 or the ISG15-USP18 complex were measured. Data were fit using a one-site binding model (also see Fig. 1c, d). **g,** Purification of the ISG15 inhibitory complex (IC) visualised by an SDS-PAGE Coomassie-stained gel (∼4 µg of the complex was loaded onto the gel). **h,** A selection of representative 2-dimensional (2D) classes used in the 3D reconstruction of the ISG15 IC. **i,** Original cryo-EM DeepEMhancer density (top panel) and segmented DeepEMhancer density (bottom panel) of the ISG15 IC in which the AlphaFold model was fit into the map. Binding is shown to occur between STAT2-USP18 and ISG15-USP18. Only the C-terminal ubiquitin-like domain (C-lobe) of ISG15 is visible.

**Supplementary Figure 3.**
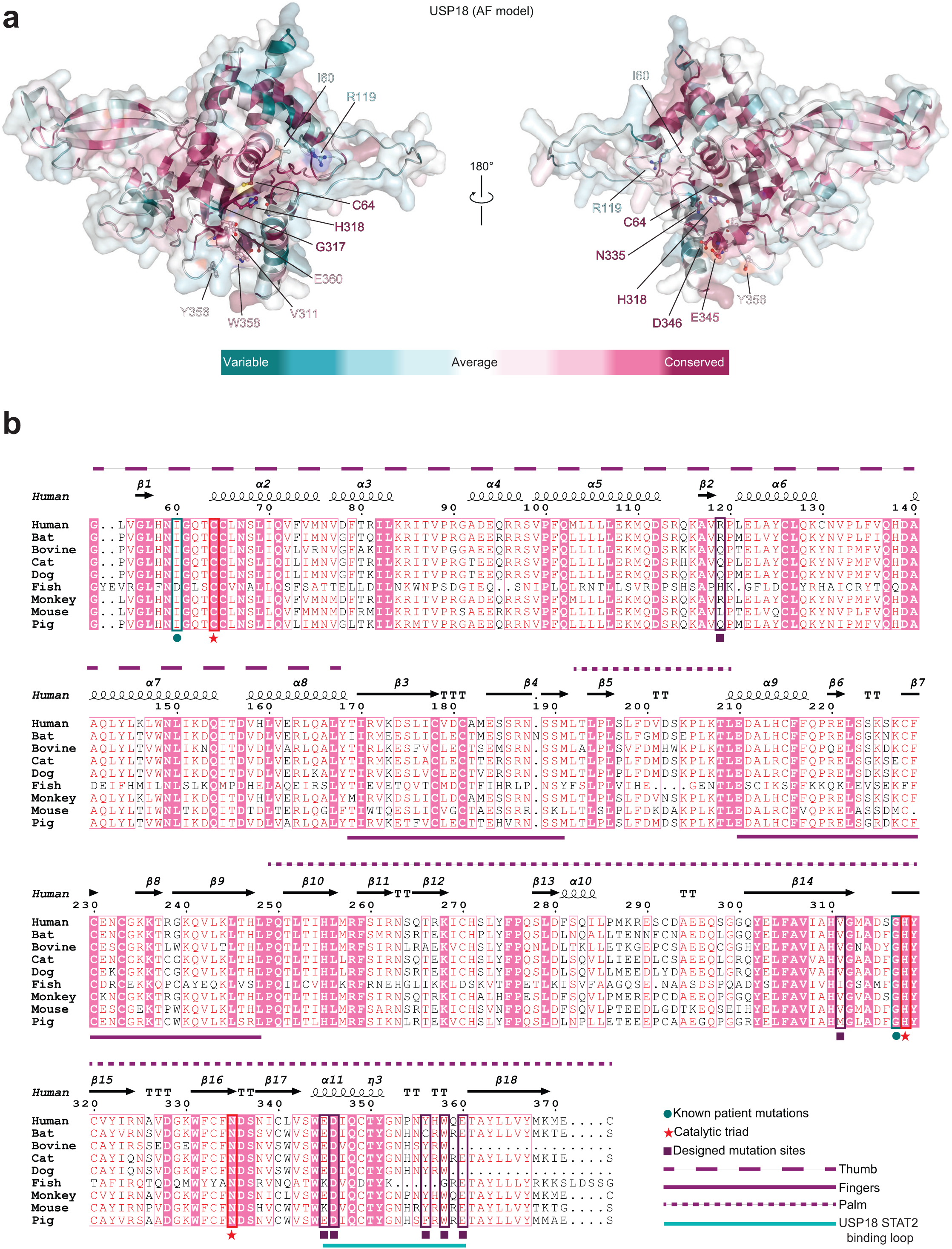
Conservation analysis of USP18 residues predicted to contact STAT2. **a**, Analysis of USP18 residue conservation was determined by ConSurf across 150 species and mapped onto the human USP18 AlphaFold model^59^. **b,** Multi-sequence alignment of USP18 from different species. Sites of patient mutations, active site residues, and mutated residues in this study are highlighted and represented with teal circles, red stars, and purple squares, respectively. T-Coffee and ESPript were used to perform the sequence alignments^57,58^.

**Supplementary Figure 4.**
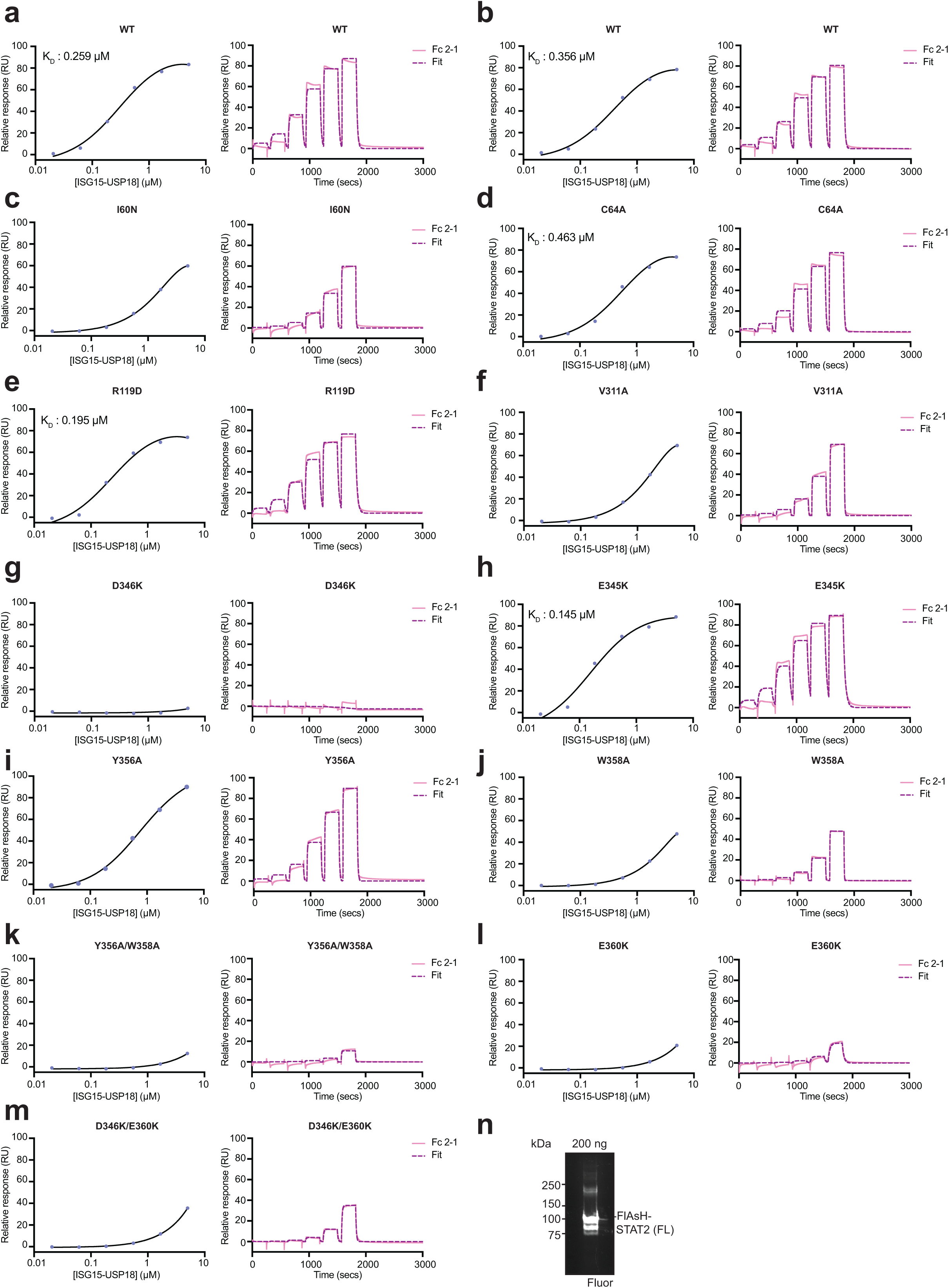
SPR binding assays of ISG15-USP18 mutants against wild-type STAT2. **a-m**, A representative example of three independent binding experiments is shown for ISG15-USP18 mutants against wild-type STAT2 (n=3). The ISG15-USP18 complexes analysed were as follows: **a, b,** wild-type (WT) **c,** I60N **d,** C64A **e,** R119D **f,** V311A **g,** D346K **h,** E345K **i,** Y356A **j,** W358A **k,** Y356A/W358A **l,** E360K **m,** D346K/E360K. For all experiments, wild-type STAT2 was immobilised onto a Strep-Tactin coupled sensor chip and association and dissociation rates of wild-type or mutant ISG15-USP18 were determined. Data were fit using a one-site binding model (also see Fig. 3d). **n,** Lumio Green labelled FlAsH-STAT2 was loaded onto an SDS-PAGE gel (200 ng) and visualised by fluorescence imaging (Fluor).

**Supplementary Figure 5.**
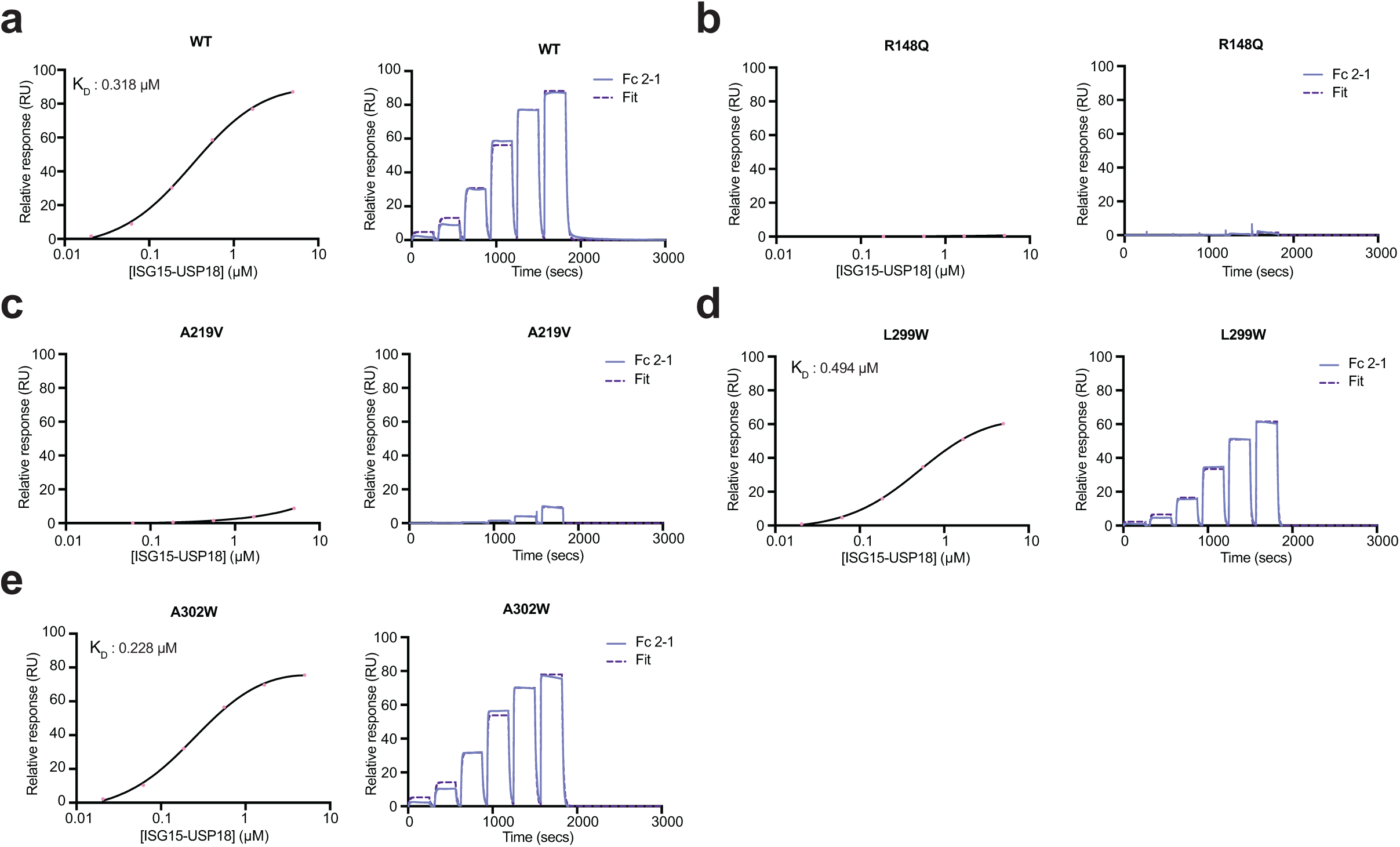
SPR binding assays of wild-type ISG15-USP18 against STAT2 mutants. **a-e**, A representative example of three independent binding experiments is shown for wild-type ISG15-USP18 against STAT2 mutants (n=3). STAT2 mutants used for these experiments were as follows: **a,** wild-type (WT) **b,** R148Q **c,** A219V **d,** L299W **e,** A302W. For each experiment wild-type and each STAT2 mutant were immobilised onto a Strep-Tactin coupled sensor chip and association and dissociation rates of wild-type ISG15-USP18 were determined. Data were fit using a one-site binding model (also see Fig 4c).

**Supplementary Table 1.** Summary of the binding parameters for the ISG15-USP18 mutant STAT2 interactions. Errors represent the standard deviation from the mean of three independent replicates (n=3). ND = not determined. For k_on_ and k_off_ values the exponential notation also applies to the standard deviation values.

| Immobilised ligand | Analyte | KD 1:1 ($\mu\text{M}$) | $k_{\text{on}}$ ($\text{M}^{-1}\text{s}^{-1}$) | $k_{\text{off}}$ ($\text{s}^{-1}$) | $K_{\text{D}}$ SS ($\mu\text{M}$) |
| --- | --- | --- | --- | --- | --- |
| STAT2<sub>wt</sub> | ISG15-USP18<sub>wt</sub> | $0.420 \pm 0.044$ | $1.48\text{E}+05 \pm 0.09$ | $6.17\text{E}-02 \pm 0.46$ | $0.366 \pm 0.065$ |
| STAT2<sub>wt</sub> | ISG15-USP18<sub>I60N</sub> | ND | ND | ND | ND |
| STAT2<sub>wt</sub> | ISG15-USP18<sub>C64A</sub> | $0.587 \pm 0.009$ | $1.18\text{E}+05 \pm 0.04$ | $6.92\text{E}-02 \pm 0.29$ | $0.467 \pm 0.008$ |
| STAT2<sub>wt</sub> | ISG15-USP18<sub>R119D</sub> | $0.336 \pm 0.028$ | $1.05\text{E}+05 \pm 0.10$ | $3.60\text{E}-02 \pm 0.15$ | $0.206 \pm 0.013$ |
| STAT2<sub>wt</sub> | ISG15-USP18<sub>V311A</sub> | ND | ND | ND | ND |
| STAT2<sub>wt</sub> | ISG15-USP18<sub>D346K</sub> | ND | ND | ND | ND |
| STAT2<sub>wt</sub> | ISG15-USP18<sub>E345K</sub> | $0.276 \pm 0.039$ | $1.09\text{E}+05 \pm 0.14$ | $2.98\text{E}-02 \pm 0.08$ | $0.175 \pm 0.030$ |
| STAT2<sub>wt</sub> | ISG15-USP18<sub>Y356A</sub> | ND | ND | ND | ND |
| STAT2<sub>wt</sub> | ISG15-USP18<sub>W358A</sub> | ND | ND | ND | ND |
| STAT2<sub>wt</sub> | ISG15-USP18<sub>Y356A+W358A</sub> | ND | ND | ND | ND |
| STAT2<sub>wt</sub> | ISG15-USP18<sub>E360K</sub> | ND | ND | ND | ND |
| STAT2<sub>wt</sub> | ISG15-USP18<sub>D346K+E360K</sub> | ND | ND | ND | ND |

ND: Not determined

**Supplementary Table 2.** Summary of the binding parameters for the STAT2 mutant ISG15-USP18 interactions. Errors represent the standard deviation from the mean of three independent replicates (n=3). ND = not determined. For k_on_ and k_off_ values the exponential notation also applies to the standard deviation values.

| Immobilised ligand | Analyte | KD 1:1 (μM) | k<sub>on</sub> (M<sup>-1</sup>s<sup>-1</sup>) | k<sub>off</sub> (s<sup>-1</sup>) | K<sub>D</sub> SS (μM) |
| --- | --- | --- | --- | --- | --- |
| STAT2<sub>wt</sub> | ISG15-USP18<sub>wt</sub> | 0.383 ± 0.004 | 1.47E+05 ± 0.15 | 5.62E-02 ± 0.59 | 0.311 ± 0.061 |
| STAT2<sub>R148Q</sub> | ISG15-USP18<sub>wt</sub> | ND | ND | ND | ND |
| STAT2<sub>A219V</sub> | ISG15-USP18<sub>wt</sub> | ND | ND | ND | ND |
| STAT2<sub>L299W</sub> | ISG15-USP18<sub>wt</sub> | 0.555 ± 0.037 | 1.48E+05 ± 0.32 | 8.53E-02 ± 1.34 | 0.459 ± 0.035 |
| STAT2<sub>A302W</sub> | ISG15-USP18<sub>wt</sub> | 0.303 ± 0.009 | 1.87E+05 ± 0.29 | 5.69E-02 ± 0.74 | 0.230 ± 0.004 |

ND: Not determined

## Methods

### Molecular biology

The following genes were cloned into the pLIB baculovirus-insect cell expression vector using the Gibson assembly method: human ISG15 (1-157 aa), human USP18 (1-372 aa, C-terminal 6xHis tag), human ΔN-USP18 (36-372 aa, C-terminal 6xHis tag), human USP18 (49-372 aa, N-terminal GST tag) and human STAT2 (1-851 aa, C-terminal Twin-Strep tag). Full-length Zika virus (ZIKV) NS5 (1-903 aa, N-terminal His-SUMO tag) was cloned into the pCoofy5 bacterial expression vector. Briefly, vectors containing the appropriate tags were first linearised and then purified by DpnI digestion and gel extraction. Gene inserts were PCR amplified with overlapping vector specific DNA sequences and purified by DpnI digestion and gel extraction. The gene inserts (50 ng) and linearised vectors (25 ng) were added to Gibson assembly mix (1:1 vol/vol) and the reaction was performed at 50**°**C for one hour. For vectors containing multigene assemblies (i.e., biGBac assembly) methods by Weissmann et al., were used^37^. Patient and structural mutations were introduced into the pLIB vectors via quick change PCR.

### Insect cell tissue culture

Sf9 cells (Thermo Fisher, Cat. no. 11496015) and High Five cells (Cat. no. B85502) were cultured in SF900 II and ESF 921 media supplemented with antibiotic-antimycotic (Gibco), respectively. To prepare baculovirus stocks, bacmid DNA was transfected into adherent SF9 cells (2 × 10^6^ cells/35 mm well) using the FuGENE HD transfection reagent (Promega) and incubated for five days at 27**°**C. The baculovirus was then harvested by collecting the media from the infected cells. The virus was amplified by adding 500 µL of the baculovirus to adherent SF9 cells (0.8 × 10^6^ cells/15 cm dish) and incubated for three days at 27**°**C before the media was harvested. This same protocol was repeated to generate baculovirus for large-scale protein expression. For protein expression, High Five suspension cells grown to a density of ∼2 million cells/mL were infected with baculovirus prepared from Sf9 cells. The infection was left to incubate for three days at 27**°**C. Cells were then harvested, flash frozen, and stored at -80**°**C.

### Protein purification

All proteins except for ZIKV NS5 were expressed and purified from High Five cells using the baculovirus expression system. Prior to sonication, all lysis buffers were supplemented with the following protease inhibitors and nuclease unless indicated otherwise: 10 µg/mL leupeptin, 20 µg/mL aprotinin, 1 mM PMSF, 20-40 units Benzonase (Merck) and 1 tablet of EDTA-free complete protease inhibitor (Roche).

#### ISG15 IC

Cell pellets for the ISG15-USP18-STAT2 ternary complex were resuspended in lysis buffer (50 mM Tris pH 8.0, 150 mM NaCl, 2 mM β-mercaptoethanol). Suspended cells were then sonicated, centrifuged, and the supernatant was incubated with Ni-NTA agarose (Abcam). Protein was eluted from the resin with lysis buffer supplemented with 250 mM imidazole. The eluted protein was then incubated with Strep-Tactin XT 4Flow resin (IBA-Lifesciences) before elution with 1X BXT biotin elution buffer (IBA-Lifesciences). Eluted protein was then buffer exchanged into storage buffer (50 mM Tris pH 8.0, 150 mM NaCl, 2 mM DTT) using a PD10 column (Cytivia). To further assess complex stability high-performance liquid chromatography (HPLC) was performed on a Vanquish instrument (Thermo Fisher) with an Advance Bio SEC 300Å 2.7 µM 4.6×300 mm column (Agilent). 15 µL of 26 µM ISG15 IC was loaded onto the column. Fractions containing protein were then visualised by SDS-PAGE and silver staining.

#### Human STAT2

Wild-type STAT2 (1-851aa), STAT2 mutants, and FlAsH-STAT2 cell pellets were resuspended in lysis buffer (50 mM Tris pH 8.0, 150 mM NaCl, 2 mM β-mercaptoethanol). Suspended cells were then sonicated, centrifuged, and the supernatant was incubated with Strep-Tactin XT 4Flow resin (IBA-Lifesciences). Bound protein was then eluted in 1X BXT biotin elution buffer (IBA-Lifesciences) and further purified using ion exchange (ResourceQ column) and size exclusion chromatography (Superdex 200 increase 10/300 GL). Purified proteins were kept in storage buffer (50 mM Tris pH 8.0, 150 mM NaCl, 2 mM DTT) at -80**°**C.

#### ISG15-USP18 complex

ISG15-USP18 wild-type and mutant cell pellets were resuspended in lysis buffer (50 mM Tris pH 8.8, 300 mM NaCl, 5% glycerol, 15 mM imidazole, 2 mM β-mercaptoethanol). Suspended cells were then sonicated, centrifuged, and the supernatant was incubated with Ni-NTA agarose (Abcam). Protein was eluted with lysis buffer supplemented with 250 mM imidazole. Eluted protein was further purified using size exclusion chromatography (Superdex 75 increase 10/200 GL). Purified proteins were kept in storage buffer (50 mM Tris pH 8.0, 300 mM NaCl, 2 mM DTT) at -80**°**C.

#### Human USP18

N-terminally GST tagged human USP18 (49-372 aa) was expressed and purified as previously described^18^. Briefly, Human USP18 pellets were sonicated, centrifuged, and the supernatant was incubated with Glutathione Sepharose 4B (GE Healthcare). Bound protein was then eluted with lysis buffer supplemented with 10 mM glutathione and further purified using ion exchange (ResourceQ column) and gel filtration (Superdex 75 increase 10/300 GL). Purified protein was kept in storage buffer (50 mM Tris pH 8.8, 150 mM NaCl, 2 mM DTT) at -80**°**C.

#### Zika virus NS5

Full-length NS5 was expressed as previously described^60^. Briefly, cell pellets were resuspended in lysis buffer (25 mM Tris pH 7.5, 500 mM NaCl, 20 mM imidazole, 5% glycerol, 5 mM β-mercaptoethanol) supplemented with EDTA-free complete protease inhibitor tablets (Roche), DNase, and lysozyme. Suspended cells were then sonicated, centrifuged, and the supernatant incubated with Ni-NTA agarose (Abcam). Protein was eluted with lysis buffer supplemented with 350 mM imidazole. Eluted protein was then treated with 0.2 mg of the SUMO protease SENP1 and dialysed overnight (3500 MWCO snakeskin tubing, Thermo Fisher) in lysis buffer minus imidazole. After overnight SENP1 cleavage, the protein was incubated with nickel resin to remove the cleaved His-SUMO tag. The flow through was then further purified using size exclusion chromatography (Superdex 200 increase 10/300 GL). Purified protein was kept in storage buffer (25 mM Tris pH 7.5, 500 mM NaCl, 5% glycerol and 5 mM DTT) at - 80**°**C.

### Surface plasmon resonance binding studies

All surface plasmon resonance (SPR) assays were performed on a Biacore 8K+ instrument (Cytiva). Strep-Tactin XT (IBA-Lifesciences) was immobilised across all channels of a CM5 sensor chip (Cytiva) via standard amine coupling chemistry (1:1 EDC/NHS) at pH 7.5. Wild-type and mutant STAT2 variants were diluted in SPR running buffer (50 mM Tris pH 8.0, 150 mM NaCl, 0.05% Tween-20, 2 mM DTT) and captured onto the functionalised surface to a target density of ∼100 response units (RU).

#### Single-cycle analysis

The ISG15-USP18 analyte was prepared in running buffer as a six-point, three-fold (1:3) serial dilution, starting from 5 µM down to 0.02 µM. The analyte was injected at a flow rate of 10 µL/min with an association time of 250 s and a dissociation time of 1200 s. SPR sensorgrams were recorded by relative response (RU) versus time (s). All data were analysed using the Biacore Insight Evaluation software. To determine binding kinetics, sensorgrams were globally fitted to a 1:1 kinetic binding model to derive the association (k_on_) and dissociation (k_off_) rate constants, from which the kinetic equilibrium dissociation constant (KD) was calculated. Additionally, steady-state K_D_ values were determined by plotting the maximal equilibrium response units (RU) against analyte concentration fitted to a steady-state affinity model. Both kinetic and steady-state parameters are reported only for interactions where binding could be confidently determined; mutant variants lacking a detectable response or exhibiting binding too weak to yield a reliable mathematical fit were designated as exhibiting no binding (NB).

### Cryo-EM

#### Sample preparation

For cryo-EM sample preparation 3.5 µL of 0.5 mg/mL ISG15 IC was applied to a plasma cleaned Quantifoil Cu grid (1.2/1.3, 400 mesh). Grids were then blotted using the following settings before being plunge-frozen in liquid ethane using a Vitrobot Mark VI (Thermo Fisher): humidity 100%, blotting time 3.5 s, blot force 3, wait time 6 s, drain time 2 s. Grids were then stored in liquid nitrogen.

#### Data Acquisition

Immediately prior to cryo-EM analysis the grids were clipped and placed into the microscope’s grid autoloader. Samples were then screened, and a dataset was collected using a 200 kV Glacios cryo-transmission electron microscope (cryo-TEM) equipped with a Falcon4i Direct Electron Detector (Thermo Fisher). In total 2,568 movies were recorded at a pixel size of 1.2 Å, total dose of 29 e/Å^2^, and an exposure time of 2.8 sec, with the defocus range spanning from −3.3 to −1.8 µM.

#### Data processing

CryoSPARC was used to process the dataset and reconstruct the 3D map of the ISG15–USP18–STAT2 complex. Movies were first imported into CryoSPARC with an EER upscaling factor of 2 and the EER frames were grouped into 38 movie fractions. Motion correction was performed using patch motion correction, and defocus estimation was carried out using CTFFIND4. Particles were then picked using the Blob Picker and extracted with a box size of 96 Å. A few rounds of 2D classification were performed to select the best particle stack. An initial model was generated using the ab initio reconstruction routine in CryoSPARC and was subsequently refined using homogeneous and non-uniform refinement protocols. To identify conformationally homogeneous particles, 3D classification was performed, followed by further refinement. Post-processing of the final cryo-EM map was carried out using DeepEMhancer^61^.

#### Model fitting

The AlphaFold model of the ISG15-USP18-STAT2 ternary complex was generated and rigid body fit into the DeepEMhancer density in ChimeraX. The DeepEMhancer map was segmented corresponding to each component of the complex and coloured using volume tools in ChimeraX.

### FlAsH-STAT2 fluorescence labelling

FlAsH-STAT2 was purified as described above for wild-type STAT2 with slight modifications. Size exclusion chromatography was performed into FlAsH buffer (50 mM Tris pH 8.0, 150 mM NaCl, 1 mM β-mercaptoethanol). Labelling was performed following methods by Wauer et al.,^62^. After overnight incubation, labelled protein was buffer exchanged using a Zeba column (Thermo Fisher) into FlAsH buffer with a final concentration of 2 mM β-mercaptoethanol and concentrated to ∼40 µM.

### Fluorescence polarisation binding assays

All fluorescence polarisation binding assays were performed in black flat-bottomed small volume 384-well plates (Thermo Fisher). Fluorescence polarisation measurements were made with a 482 nM excitation filter and a 530 nM emission filter at a target gain of 300 using a Clariostar Plus plate reader (BMG Labtech).

#### End-point binding assays

FlAsH-STAT2 was diluted to a 2x concentration of 200 nM into assay buffer (50 mM Tris pH 8.0, 150 mM NaCl, 2 mM DTT). For assays in Fig. 2e, free GST, GST-USP30 (DU36294, https://mrcppureagents.dundee.ac.uk/), and ISG15-USP18 were diluted to 2x concentrations of 0.5 µM, 1 µM and 2 µM. For assays in Fig. 3c, ISG15-USP18 wild-type and mutants were diluted to a 2x concentration of 1 µM. FlAsH-STAT2 and each binding partner were mixed at a 1:1 ratio (vol/vol) and the plate was subsequently incubated for 15 minutes prior to fluorescent polarisation measurements. 200 flashes per well were performed every 0.2 seconds. Measurements were performed in triplicate and error bars represent the standard deviation from the mean. Data was analysed using GraphPad Prism (version 10.6.1).

##### Real-time fluorescence polarisation binding assays

Fluorescence polarisation assays were adapted from Wallace et al., with modifications as outlined below^48^. FlAsH-STAT2 was prepared at an initial 1.33x concentration of 133 nM in assay buffer. ISG15-USP18 (wild-type and Y356A/W358A) was prepared to an initial 8x concentration (12 µM), and the viral proteins (NS5 full-length, NS1B WT and NS1B-AA) were prepared at an initial 8x concentration (24 µM). 2.5 µL of protein or buffer were added to the FlAsH-STAT2 at the indicated cycle number. 200 flashes per well were performed every 70 seconds for 30 cycles. Data was analysed using GraphPad Prism (version 10.6.1).

### USP18 cleavage assays

For time-dependent proISG15 cleavage assays in Fig. 5a, 10 µM proISG15 was incubated with and without 2 µM STAT2 and 1 µM GST-tagged USP18 (49-372 aa) in assay buffer (50 mM pH 8.0, 150 mM NaCl, 2 mM DTT) for 120 minutes at 37**°**C. The reaction was quenched with LDS sample buffer at the indicated time points before being visualised by SDS-PAGE and Coomassie staining. For concentration-dependent cleavage assays in Fig. 5b, the same USP18 and proISG15 concentrations were used with increasing STAT2 concentrations (0, 1, 2, 3, 4 and 5 µM). Reactions were performed at 37**°**C for 60 minutes before being quenched and visualised by Coomassie-stained SDS-PAGE gels.

### *In silico* ISG15 IC modelling

The ISG15 IC model was generated using AlphaFold3 with the following UniProt sequences, ISG15 (UniProt: P05161, 1-157 aa), USP18 (UniProt: Q9UMW8, 36-372 aa) and STAT2 (UniProt: P52630, 1-681 aa)^39^. For the apo USP18 and ISG15-USP18 models the same sequences were input into AlphaFold3. All components were coloured by domain and structural analysis was performed using PyMOL2.

## Data availability

All data is available upon request from the corresponding author.

## Notes

### Competing Interest Statement

The authors have declared no competing interest.

## References

1. Chen, R. et al. Pattern recognition receptors: function, regulation and therapeutic potential. Signal Transduct. Target. Ther. 10, 216 (2025).

2. Ivashkiv, L. B. & Donlin, L. T. Regulation of type I interferon responses. Nat Rev Immunol 14, 36–49 (2014).

3. Philips, R. L. et al. The JAK-STAT pathway at 30: Much learned, much more to do. Cell 185, 3857–3876 (2022).

4. Schneider, W. M., Chevillotte, M. D. & Rice, C. M. Interferon-Stimulated Genes: A Complex Web of Host Defenses. Annu Rev Immunol 32, 513–545 (2014).

5. Arimoto, K.-I., Miyauchi, S., Stoner, S. A., Fan, J.-B. & Zhang, D.-E. Negative regulation of type I IFN signaling. J. Leukoc. Biol. 103, 1099–1116 (2018).

6. Haas, A. L., Ahrens, P., Bright, P. M. & Ankel, H. Interferon induces a 15-kilodalton protein exhibiting marked homology to ubiquitin. J Biol Chem 262, 11315–11323 (1987).

7. Farrell, P. J., Broeze, R. J. & Lengyel, P. Accumulation of an mRNA and protein in interferon-treated Ehrlich ascites tumour cells. Nature 279, 523–525 (1979).

8. Yuan, W. & Krug, R. M. Influenza B virus NS1 protein inhibits conjugation of the interferon (IFN)-induced ubiquitin-like ISG15 protein. Embo J 20, 362–371 (2001).

9. Zhao, C. et al. The UbcH8 ubiquitin E2 enzyme is also the E2 enzyme for ISG15, an IFN-α/β-induced ubiquitin-like protein. P Natl Acad Sci Usa 101, 7578–7582 (2004).

10. Dastur, A., Beaudenon, S., Kelley, M., Krug, R. M. & Huibregtse, J. M. Herc5, an Interferon-induced HECT E3 Enzyme, Is Required for Conjugation of ISG15 in Human Cells*. J. Biol. Chem. 281, 4334–4338 (2006).

11. Wong, J. J. Y., Pung, Y. F., Sze, N. S.-K. & Chin, K.-C. HERC5 is an IFN-induced HECT-type E3 protein ligase that mediates type I IFN-induced ISGylation of protein targets. Proc National Acad Sci 103, 10735–10740 (2006).

12. Durfee, L. A., Kelley, M. L. & Huibregtse, J. M. The Basis for Selective E1-E2 Interactions in the ISG15 Conjugation System*. J Biol Chem 283, 23895–23902 (2008).

13. Durfee, L. A., Lyon, N., Seo, K. & Huibregtse, J. M. The ISG15 Conjugation System Broadly Targets Newly Synthesized Proteins: Implications for the Antiviral Function of ISG15. Mol. Cell 38, 722–732 (2010).

14. Malakhov, M. P., Malakhova, O. A., Kim, K. I., Ritchie, K. J. & Zhang, D.-E. UBP43 (USP18) Specifically Removes ISG15 from Conjugated Proteins*. J. Biol. Chem. 277, 9976–9981 (2002).

15. Basters, A. et al. Structural basis of the specificity of USP18 toward ISG15. Nat Struct Mol Biol 24, 270–278 (2017).

16. Malakhova, O. A. et al. UBP43 is a novel regulator of interferon signaling independent of its ISG15 isopeptidase activity. Embo J 25, 2358–2367 (2006).

17. Zhang, X. et al. Human intracellular ISG15 prevents interferon-α/β over-amplification and auto-inflammation. Nature 517, 89–93 (2015).

18. Vasou, A. et al. ISG15-Dependent Stabilisation of USP18 Is Necessary but Not Sufficient to Regulate Type I Interferon Signalling in Humans. Eur. J. Immunol. 55, e202451651 (2025).

19. Arimoto, K. et al. STAT2 is an essential adaptor in USP18-mediated suppression of type I interferon signaling. Nat Struct Mol Biol 24, 279–289 (2017).

20. Perng, Y.-C. & Lenschow, D. J. ISG15 in antiviral immunity and beyond. Nat Rev Microbiol 16, 423–439 (2018).

21. Wilmes, S. et al. Receptor dimerization dynamics as a regulatory valve for plasticity of type I interferon signaling. J. Cell Biol. 209, 579–593 (2015).

22. Martin-Fernandez, M. et al. Systemic Type I IFN Inflammation in Human ISG15 Deficiency Leads to Necrotizing Skin Lesions. Cell Rep. 31, 107633 (2020).

23. Buda, G. et al. Inflammatory cutaneous lesions and pulmonary manifestations in a new patient with autosomal recessive ISG15 deficiency case report. *Allergy*, Asthma Clin. Immunol. 16, 77 (2020).

24. Al-Mayouf, S. M., Akbar, L., AlEnazi, A. & Al-Mousa, H. Autosomal Recessive ISG15 Deficiency Underlies Type I Interferonopathy with Systemic Lupus Erythematosus and Inflammatory Myositis. J. Clin. Immunol. 41, 1361–1364 (2021).

25. Burleigh, A. et al. Case Report: ISG15 deficiency caused by novel variants in two families and effective treatment with Janus kinase inhibition. Front. Immunol. 14, 1287258 (2023).

26. Napoleao, S. M. da S., et al. First Brazilian Case Report of Unrelated Patients with Identical ISG15 Mutation. J. Clin. Immunol. 45, 21 (2025).

27. Sazeides, C., et al. Human ISG15 deficiency unveils impaired healing of ulcerations via type I interferon–mediated fibrosis. J. Hum. Immun. 2, e20250011 (2026).

28. Bogunovic, D. et al. Mycobacterial Disease and Impaired IFN-γ Immunity in Humans with Inherited ISG15 Deficiency. Science 337, 1684–1688 (2012).

29. Speer, S. D. et al. ISG15 deficiency and increased viral resistance in humans but not mice. Nat Commun 7, 11496 (2016).

30. Meuwissen, M. E. C. et al. Human USP18 deficiency underlies type 1 interferonopathy leading to severe pseudo-TORCH syndrome. J. Exp. Med. 213, 1163–1174 (2016).

31. Martin-Fernandez, M. et al. A partial form of inherited human USP18 deficiency underlies infection and inflammation. J Exp Med 219, e20211273 (2022).

32. Alsohime, F. et al. JAK Inhibitor Therapy in a Child with Inherited USP18 Deficiency. New Engl J Med 382, 256–265 (2020).

33. Sun, X. et al. Novel USP18 mutations lead to severe interferonopathy responsive to JAK inhibitor. Front. Immunol. 16, 1646996 (2025).

34. Bucciol, G. et al. Human inherited complete STAT2 deficiency underlies inflammatory viral diseases. J. Clin. Investig. 133, e168321 (2023).

35. Zhu, G. et al. Type I Interferonopathy due to a Homozygous Loss-of-Inhibitory Function Mutation in STAT2. J. Clin. Immunol. 43, 808–818 (2023).

36. Gruber, C. et al. Homozygous STAT2 gain-of-function mutation by loss of USP18 activity in a patient with type I interferonopathy. J Exp Med 217, e20192319 (2020).

37. Weissmann, F., et al. biGBac enables rapid gene assembly for the expression of large multisubunit protein complexes. Proc National Acad Sci 113, E2564–E2569 (2016).

38. Burkart, C., Fan, J.-B. & Zhang, D.-E. Two Independent Mechanisms Promote Expression of an N-terminal Truncated USP18 Isoform with Higher DeISGylation Activity in the Nucleus*. J. Biol. Chem. 287, 4883–4893 (2012).

39. Abramson, J. et al. Accurate structure prediction of biomolecular interactions with AlphaFold 3. Nature 630, 493–500 (2024).

40. Kazi, N. H., Klink, N., Gallant, K., Kipka, G.-M. & Gersch, M. Chimeric deubiquitinase engineering reveals structural basis for specific inhibition of the mitophagy regulator USP30. Nat. Struct. Mol. Biol. 32, 1776–1786 (2025).

41. Espada, C. E. et al. ISG15/USP18/STAT2 is a molecular hub regulating IFN I-mediated control of Dengue and Zika virus replication. Front. Immunol. 15, 1331731 (2024).

42. Jové, V. et al. Type I interferon regulation by USP18 is a key vulnerability in cancer. iScience 27, 109593 (2024).

43. Duncan, C. J. A. et al. Severe type I interferonopathy and unrestrained interferon signaling due to a homozygous germline mutation in STAT2. Sci. Immunol. 4, (2019).

44. Sarkar, L., Liu, G. & Gack, M. U. ISG15: its roles in SARS-CoV-2 and other viral infections. Trends Microbiol. 31, 1262–1275 (2023).

45. Munnur, D., Banducci-Karp, A. & Sanyal, S. ISG15 driven cellular responses to virus infection. Biochem. Soc. Trans. 50, 1837–1846 (2022).

46. Miorin, L., Maestre, A. M., Fernandez-Sesma, A. & García-Sastre, A. Antagonism of type I interferon by flaviviruses. Biochem. Biophys. Res. Commun. 492, 587–596 (2017).

47. Parks, G. D. & Alexander-Miller, M. A. Paramyxovirus Activation and Inhibition of Innate Immune Responses. J. Mol. Biol. 425, 4872–4892 (2013).

48. Wallace, I. et al. Insights into the ISG15 transfer cascade by the UBE1L activating enzyme. Nat. Commun. 14, 7970 (2023).

49. Franklin, T. G. & Pruneda, J. N. A High-Throughput Assay for Monitoring Ubiquitination in Real Time. Front. Chem. 7, 816 (2019).

50. Wang, B. et al. Structural basis for STAT2 suppression by flavivirus NS5. Nat. Struct. Mol. Biol. 27, 875–885 (2020).

51. Huynh, K. W. et al. Insight into the scaffolding function of USP18 from a high resolution cryo-EM structure of STAT2-USP18-ISG15 ternary complex. bioRxiv 2026.02.12.705587 (2026) doi:10.64898/2026.02.12.705587.

52. Grant, A. et al. Zika Virus Targets Human STAT2 to Inhibit Type I Interferon Signaling. Cell Host Microbe 19, 882–890 (2016).

53. Ren, W., et al. Zika virus NS5 protein inhibits type I interferon signaling via CRL3 E3 ubiquitin ligase-mediated degradation of STAT2. Proc. Natl. Acad. Sci. 121, e2403235121 (2024).

54. Fernández, D. J., Hess, S. & Knobeloch, K.-P. Strategies to Target ISG15 and USP18 Toward Therapeutic Applications. Front. Chem. 7, 923 (2020).

55. Hess, S. et al. A NanoBRET-based assay monitoring interactions in the USP18 signaling hub identifies the first cell-penetrant small molecule compromising USP18/ISG15 binding. bioRxiv 2025.03.30.646166 (2025) doi:10.1101/2025.03.30.646166.

56. Waterhouse, A. M., Procter, J. B., Martin, D. M. A., Clamp, M. & Barton, G. J. Jalview Version 2—a multiple sequence alignment editor and analysis workbench. Bioinformatics 25, 1189–1191 (2009).

57. Notredame, C., Higgins, D. G. & Heringa, J. T-coffee: a novel method for fast and accurate multiple sequence alignment11Edited by J. Thornton. J Mol Biol 302, 205–217 (2000).

58. Robert, X. & Gouet, P. Deciphering key features in protein structures with the new ENDscript server. Nucleic Acids Res 42, W320–W324 (2014).

59. Landau, M. et al. ConSurf 2005: the projection of evolutionary conservation scores of residues on protein structures. Nucleic Acids Res. 33, W299–W302 (2005).

60. Wang, B. et al. The structure of Zika virus NS5 reveals a conserved domain conformation. Nat. Commun. 8, 14763 (2017).

61. Sanchez-Garcia, R. et al. DeepEMhancer: a deep learning solution for cryo-EM volume post-processing. Commun. Biol. 4, 874 (2021).

62. Wauer, T., Simicek, M., Schubert, A. & Komander, D. Mechanism of phospho-ubiquitin-induced PARKIN activation. Nature 524, 370–374 (2015).

